# A functional genomics screen identifies novel drivers of FR900359 resistance in uveal melanoma cells

**DOI:** 10.64898/2026.05.29.728309

**Authors:** Sarina D. Murray, Jesse D. Riordan, Eric R. Anderson, Michael D. Onken, Kendall J. Blumer, Christopher S. Stipp, Adam J. Dupuy

**Author notes:** co-corresponding authors: Adam J. Dupuy, Christopher S. Stipp. Co-author contacts: Sarina D. Murray Jesse D. Riordan Eric R. Anderson Michael D. Onken Kendall J. Blumer.

## Abstract

Uveal melanoma (UM) is the most common form of intraocular cancer in adults and has a median survival rate of ∼1 year after metastasis occurs. Metastatic UM is largely refractory to treatment and there are no effective pharmacological therapies, resulting in poor overall survival. Activating mutations in GNAQ and GNA11 proteins (GNAQ/11) are the oncogenic initiators in >90% UM cases. While there are no targeted therapies yet identified for the GNAQ/11 oncoproteins, a natural compound called FR900359 (FR) is a selective inhibitor for both oncogenic and wild type GNAQ/11. We performed a functional genomics screen to identify drivers of FR resistance in two UM cell lines (92.1 and MEL202). The screen identified eleven genes as candidate FR resistance drivers in both cell lines. Over-expression of five of these genes (ABCB1, PLCB4, GRM1, PLCE1, PDGFRB) was predicted to provide resistance to FR treatment. Enforced expression of ABCB1 or PLCB4 did not provide immediate resistance to FR, although over-expression of either transgene led to the emergence of resistant colonies at a much higher rate than occurs spontaneously in parental cells. We show that a relatively small fraction of UM cells can tolerate the initial over-expression of PLCB4 and ABCB1, but FR treatment leads to expansion of this cell population. Expression of an ABCB1-tGFP fusion protein was used to isolate drug naïve UM cells. We show that these cells are uniformly resistant to FR, unlike the bulk tumor cell population. Finally, additional experiment of the drug naïve ABCB1-tGFP+ UM cells led to the observation that these cells exhibit a significantly lower rate of protein translation, like BAP1-deficient UM cells. These findings suggest that resistance to targeted GNAQ/11 inhibitors is dictated by interaction between acquired genetic alterations and epigenetic states within heterogenous UM cell populations.

## Introduction

Uveal melanoma (UM) is the leading form of intraocular cancer in adults, the second most common form of melanoma, and although rare, has a median survival rate of ∼1 year after disease progression occurs (Albert, Ryan, & Borden, 1996; Bedikian et al., 1995; Diener-West et al., 2004; Einhorn, Burgess, & Gottlieb, 1974; Eskelin, Pyrhonen, Hahka-Kemppinen, Tuomaala, & Kivela, 2003; Frenkel et al., 2009; Gragoudas et al., 1991; Kim, Lane, & Gragoudas, 2010; Kodjikian et al., 2005; Lorenzo et al., 2019; Marshall et al., 2013; Nicholas et al., 2018; Piperno-Neumann et al., 2015; Rietschel et al., 2005; Rivoire et al., 2005; Willson et al., 2001). More than half of UM cases result in clinical metastases and the liver is the first site involved in over 90% of these patients (Albert et al., 1996; Bedikian et al., 1995; Diener-West et al., 2004; Einhorn et al., 1974; Eskelin et al., 2003; Frenkel et al., 2009; Gragoudas et al., 1991; Kim et al., 2010; Kodjikian et al., 2005; Lorenzo et al., 2019; Marshall et al., 2013; Nicholas et al., 2018; Piperno-Neumann et al., 2015; Rietschel et al., 2005; Rivoire et al., 2005; Willson et al., 2001). Metastatic UM patients are largely refractory to treatment, with systemic therapy showing no evidence for a longer median overall survival (OS) than the ∼10–16 months typical for untreated patients, and surgical resection providing only 6 months longer median OS for some patients (Carvajal et al., 2022; Rantala, Hernberg, Piperno-Neumann, Grossniklaus, & Kivela, 2022; Schillo et al., 2024).

Activating mutations in *GNAQ* or *GNA11* are thought to be the initiating oncogenic event in ∼90% of UM cases. These genes encode G protein alpha subunits which are regulated by the binding and hydrolysis of GTP. Mutation of codons Q209 or R183 constitutively activate these proteins by disrupting the GTPase activity, locking the proteins in an active conformation (Gupta et al., 2015; Lietman & McKean, 2022). FR900359 (FR) is a natural compound that inhibits GNAQ/11 proteins by preventing guanine nucleotide exchange (Lapadula et al., 2019; Nishimura et al., 2010; Takasaki et al., 2004; Todd et al., 2024). Oncogenic GNAQ/11 is also inhibited by FR, presumably by the same mechanism following spontaneous hydrolysis of the bound GTP, although there is some evidence that the mechanism of inhibition can be influenced by GNAQ/11 subcellular localization (Randolph et al., 2022). FR is highly selective for the GNAQ/11/14 subfamily (Schrage et al., 2015), and importantly, FR treatment of GNAQ/11 mutated UM cells induces cytostasis and/or cytotoxicity while FR-treated cutaneous melanoma cells driven by MAPK pathway alterations are unaffected (Davies et al., 2002; Hodis et al., 2012; Lapadula et al., 2019; Lietman & McKean, 2022; Onken et al., 2008; Saldanha et al., 2004; Schrage et al., 2015). Although FR has been used as a tool compound to inhibit oncogenic GNAQ/11, understanding how UM tumor cells evolve resistance to GNAQ/11 inhibition has become an important priority directly relevant to a clinical strategy targeting UM.

To achieve this goal, we performed a functional genomics screen to identify drivers of FR resistance in two UM cells lines (92.1, MEL202). This approach utilized a cell-based Sleeping Beauty (SB) transposon insertional mutagenesis screening approach that we recently developed (Feddersen, Schillo, et al., 2019; Feddersen, Wadsworth, et al., 2019; Schillo et al., 2024). The SB screen identified eleven candidate drivers of FR resistance that were independently identified in both UM cell lines. Over-expression of ABCB1 (ATP-binding cassette sub-family B member 1) or PLCB4 (phospholipase C beta 4) were the most common mechanisms observed in the FR resistance screen for both cell lines. Transgenic over-expression of ABCB1 or PLCB4 provided resistance to FR, maintaining MAPK pathway activation and allowing continued cell proliferation. Surprisingly, stable transgenic over-expression of ABCB1 and PLCB4 was tolerated in only a fraction of UM cells, but FR treatment led to the outgrowth of a more homogeneous population of cells expressing each transgene. Furthermore, we show that a prospectively isolated homogenous population of UM cells that tolerate ABCB1 over-expression display primary resistance to GNAQ/11 inhibition by FR. Collectively, these findings show that PLCB4 and ABCB1 promote resistance to GNAQ/11 inhibitors in a subset of UM cells. The observation that FR-resistance drivers function in some, but not all UM cells suggests that oncogenic GNAQ/11 signaling may not function uniformly in all UM cells. Future insights into the detailed mechanism of the FR resistance drivers identified in our study could provide key insights into the complexity of oncogenic GNAQ/11 signaling in heterogenous UM cell populations.

## Results

### Sleeping Beauty mutagenesis screen identifies FR resistance drivers in 92.1 and MEL202 cells

We first determined an optimal dose of FR900359 (FR) to use in our genetic screen by generating dose response curves for both 92.1 and MEL202 cell lines (**Figure 1a**). The rationale was to identify the FR concentration that provides a cytostatic effect on each cell line without inducing rapid cell death to maximize the number of cells that are available for selection during the genetic screen. We identified 50 nM as the optimal FR dose for 92.1 and MEL202. Next, we engineered cells to express the hyperactive SB100X transposase. Following verification of SB100X expression, transposon mutagenesis was initiated by transfection of a plasmid vector (pT2/Onc3) containing a mutagenic SB transposon (Feddersen, Wadsworth, et al., 2019). A non-mutagenized control cell population was also produced through transfection of a standard, nonintegrating EGFP expression plasmid. Both control and mutagenized cell populations were treated with FR beginning 24 hours after transfection, and the media and drug were replenished weekly until the endpoint (*i.e.,* appearance of macroscopic resistant colonies). Resistant colonies derived from each independently transfected population of cells were collected as a pooled population at the endpoint of each experiment. In total, we generated 10 populations of FR-resistant cells (*i.e.,* derived from independent pT2/Onc3 transfections) over four experiments for both 92.1 and MEL202.

**Figure 1.**
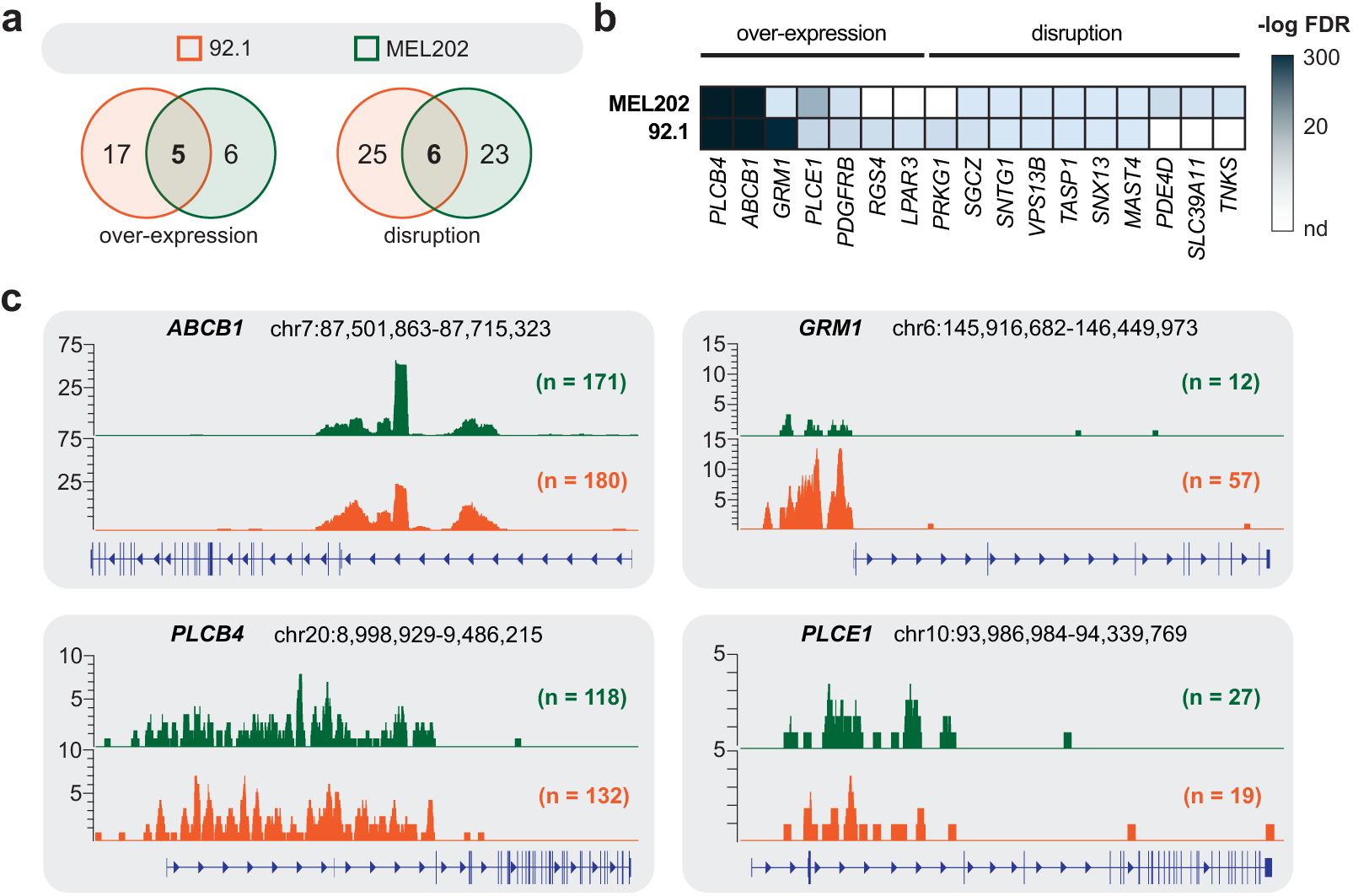
*Sleeping Beauty* mutagenesis screen to identify potential FR resistance drivers. **(a)** Analysis of screen results from 92.1 and Mel202 cell lines identified both shared and unique drivers of FR resistance. **(b)** Heatmap shows the relative differences in selected drivers of FR resistance across the two cell lines. **(c)** Transposon insertion profiles show the position and density of transposon insertions of *ABCB1, PLCB4, GRM1*, and *PLCE1*.

Transposon-induced mutations were identified essentially as previously reported (Feddersen, Wadsworth, et al., 2019). Briefly, genomic DNA was extracted from all pooled cell populations, and ∼1 µg of gDNA was fragmented to an average size of 300-800 bp. End-repair and adaptor ligation was then performed, followed by two rounds of nested PCR to amplify and barcode the transposon junction sequences. Barcoded amplicons were then pooled and sequenced on an Illumina NovaSeq-6000. Raw sequences were trimmed to remove adaptor and transposon sequences, and the remaining genomic DNA sequences were mapped to the human reference genome (GRCh38). Sequence alignments were then parsed to create a set of nonredundant transposon insertion sites within each DNA sample. These data were then filtered to eliminate low-abundance insertion events, formatted as a GFF3 file, and subjected to gene-centric common insertion site analysis (gCIS) to identify genes that exhibit a statistically significant enrichment in transposon-induced mutation associated with FR resistance (Feddersen, Wadsworth, et al., 2019). Details regarding each step can been found in the supplemental methods.

The SB mutagenesis screen identified 53 and 40 candidate drivers of FR resistance in 92.1 and MEL202 cells, respectively (**Table S1, S2**). Each driver gene was identified in at least 3 biological replicates with an FDR < 0.05. Among the benefits of SB mutagenesis is that the T2/Onc3 transposon can over-express or disrupt endogenous gene expression, depending on the context of the insertion relative to the gene (Feddersen, Wadsworth, et al., 2019). Eleven of the identified candidate resistance drivers were identified in both cell lines with the same predicted mechanism (5 over-expressed, 6 disrupted) (**Figure 1a**). Among the over-expressed candidates are two phospholipases (PLCB4, PLCE1), a drug efflux pump (ABCB1), a G-protein-coupled receptor (GRM1), and a receptor tyrosine kinase (PDGFRB). Transposon disruption is the predicted mechanism for the remaining 6 shared candidates (SGCZ, SNTG1, VPS13B, TASP1, SNX13, MAST4). As we have observed in prior drug resistance screens using SB mutagenesis (Feddersen, Schillo, et al., 2019; Schillo et al., 2024), transposon-induced over-expression tends to identify candidates with significantly lower FDR values than candidates that appear to have transposon disruption. Transposon-induced over-expression is achieved by a single transposon insertion event while gene inactivation would require transposon-induced gene disruption of all alleles, a lower probability event. The FR resistance screen shows a similar statistical trend with the strongest candidates showing transposon-induced over-expression (**Table S1, S2**).

#### Validation of ABCB1 and PLCB4 as FR resistance drivers

The SB screen identified ABCB1 and PLCB4 as the strongest drivers of FR resistance in both cell lines. PLCB4 (phospholipase C, beta 4) is a known downstream target of GNAQ/11, and activating mutations of *PLCB4* are associated with uveal melanoma and are mutually exclusive of *GNAQ*/*GNA11* mutations (Phan, Kim, Wei, Tall, & Smrcka, 2021). ABCB1 (ATP-binding cassette sub-family B member 1) is a drug-efflux pump that has been implicated in resistance to a wide variety of compounds (Skinner, Palkar, & Hong, 2023). We observed >100 independent transposon insertions within the promoter/5’end of both *PLCB4* and *ABCB1* (**Figure 1c**), and there was a clear orientation bias of the insertion events that is consistent with transposon-induced over-expression as the mechanism of FR resistance. This result provided the foundation to investigate ABCB1 and PLCB4 as drivers of FR resistance in uveal melanoma cell lines.

We generated vectors to stably express full-length coding sequences for ABCB1 (NM_001348946.2) and PLCB4 (NM_001377142.1) along with a hygromycin selection cassette. Both 92.1 and MEL202 cells were transfected with the PB-ABCB1-PGKhyg, PB-PLCB4-PGKhyg, or PB-PGKhyg (empty vector) plasmids and selected in hygromycin. Drug-resistant populations were then evaluated for FR resistance in a resazurin assay (not shown). Unexpectedly, neither transgene provided significant resistance to the cytostatic effects of FR when measured after seven days of exposure despite strong statistical evidence from the SB genetic screen.

To generate a more dynamic and quantitative assessment of each cell line’s response to FR over time, we performed an experiment using 92.1 cells in which the number of cell doublings was monitored over several weeks while cells were maintained under a chronic dose of FR (**Figure 2**). Unlike the resazurin assay, this approach allowed us to observe the emergence of FR resistance over a longer time course. As expected, FR treatment of empty vector control cells potently suppressed their proliferation and eventually caused cell death after several weeks. By comparison, drug-naïve ABCB1 transgenic cells rapidly developed resistance to FR after an initial period of drug response (**Figure 2b**). A similar trend was observed in PLCB4 transgenic cells, although the emergence of FR-resistant cells did not occur for several weeks (**Figure 2b**). Continued passaging of both ABCB1 and PLCB4 cells under constant FR selection eventually produced populations of cells with a doubling time that is indistinguishable from vehicle-treated control cells, suggesting that these cells are insensitive to FR treatment (**Figure 2c**). This experiment was repeated in the Mel202 cell line, and we observed the same trend in which ABCB1 and PLCB4 transgenic expression drove the emergence of FR resistant cells over time (**Supp. Figure 1**).

**Figure 2.**
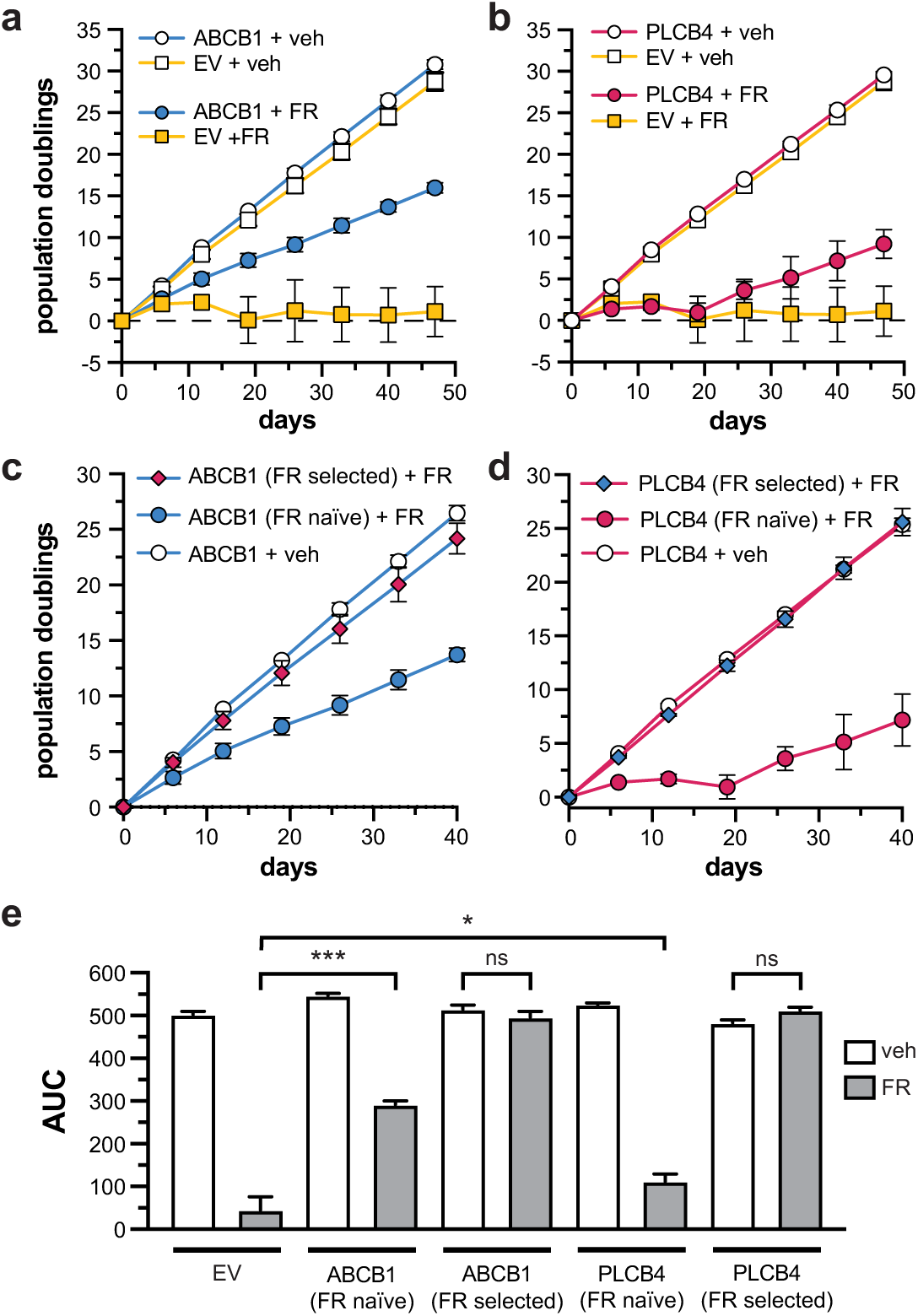
Validation of FR resistance mediated by over expression of ABCB1 and PLCB4 in 92.1 cells. Cell population doubling was monitored over several weeks in the presence of FR or vehicle for cells over expression ABCB1 (**a**) or PLCB4 (**b**). Over expression of either ABCB1 (**c**) or PLCB4 (**d**) led to the emergence of a cell populations that are indifferent to the presence of FR. (**e**) Quantification of the area under the curve (AUC) of the data shown in panels a-d.

#### GNAQ/11 downstream signaling in FR resistant UM cells expressing ABCB1 or PLCB4

Constitutive activation of GNAQ/11 in uveal melanoma has been shown to activate a complex signaling network downstream of PLC activation, including MAPK, PI3K-AKT, and YAP/TAZ pathways (**Supp. Figure 2**)(Feng et al., 2014; Saraiva, Caissie, Segal, Edelstein, & Burnier, 2005; Van Raamsdonk et al., 2009). We evaluated the activity of these pathways via western blotting in parental and transgenic cells expressing either ABCB1 or PLCB4 cells that had been selected under chronic 5 nM FR treatment (**Figure 3**). The PI3K-AKT and YAP/TAZ pathways showed variable response to FR treatment in cells expressing either ABCB1 or PLCB4 (**Supp. Figure 3**). However, we observed that both transgenes were able to maintain MAPK pathway activation marks (pRas-GRP3, pERK, p90-RSK) in both 92.1 (**Figure 3a**) and Mel202 (**Figure 3b**) cells treated with 5 nM FR.

**Figure 3.**
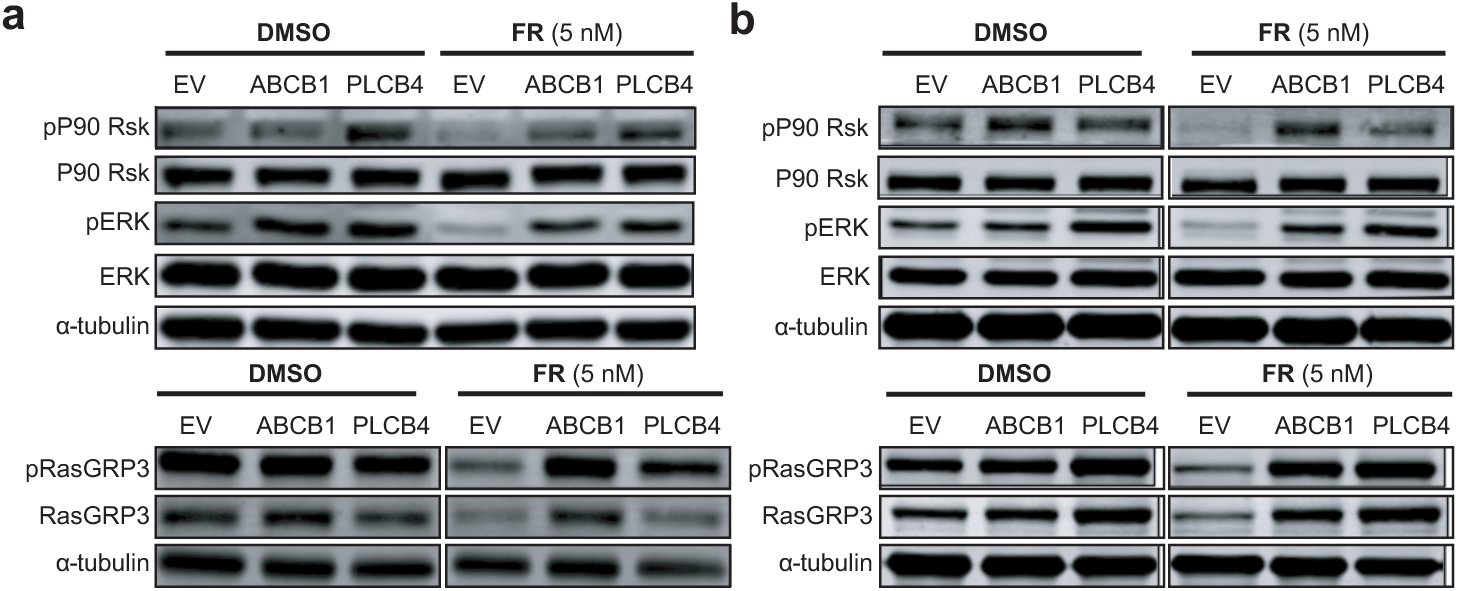
Evaluation of MAPK signaling in FR resistant cell lines. Over expression of ABCB1 or PLCB4 in 92.1 (**a**) or Mel202 (**b**) maintains MAPK signaling in the presence of FR.

#### ABCB1 and PLCB4 over-expression is tolerated in a subpopulation of UM cells

We have previously utilized SB mutagenesis to identify drivers of BRAF inhibitor resistance in cutaneous melanoma (Feddersen, Schillo, et al., 2019; Schillo et al., 2024; Zhu, Schillo, Murray, Riordan, & Dupuy, 2023). In these studies, over-expression of candidate genes produced immediate drug resistance in cells, in contrast to ABCB1 and PLCB4 which drove the emergence of drug-resistant UM cells over several passages (**Figure 2**). All transgenic cells had undergone puromycin selection to select for stable transgene integration. Western blotting had also verified elevated expression of the candidate proteins. However, FR-resistant cells had elevated expression of the transgene, suggesting that FR treatment selected for outgrowth of a subpopulation of cells with high transgene expression (**Figure 4a**).

**Figure 4.**
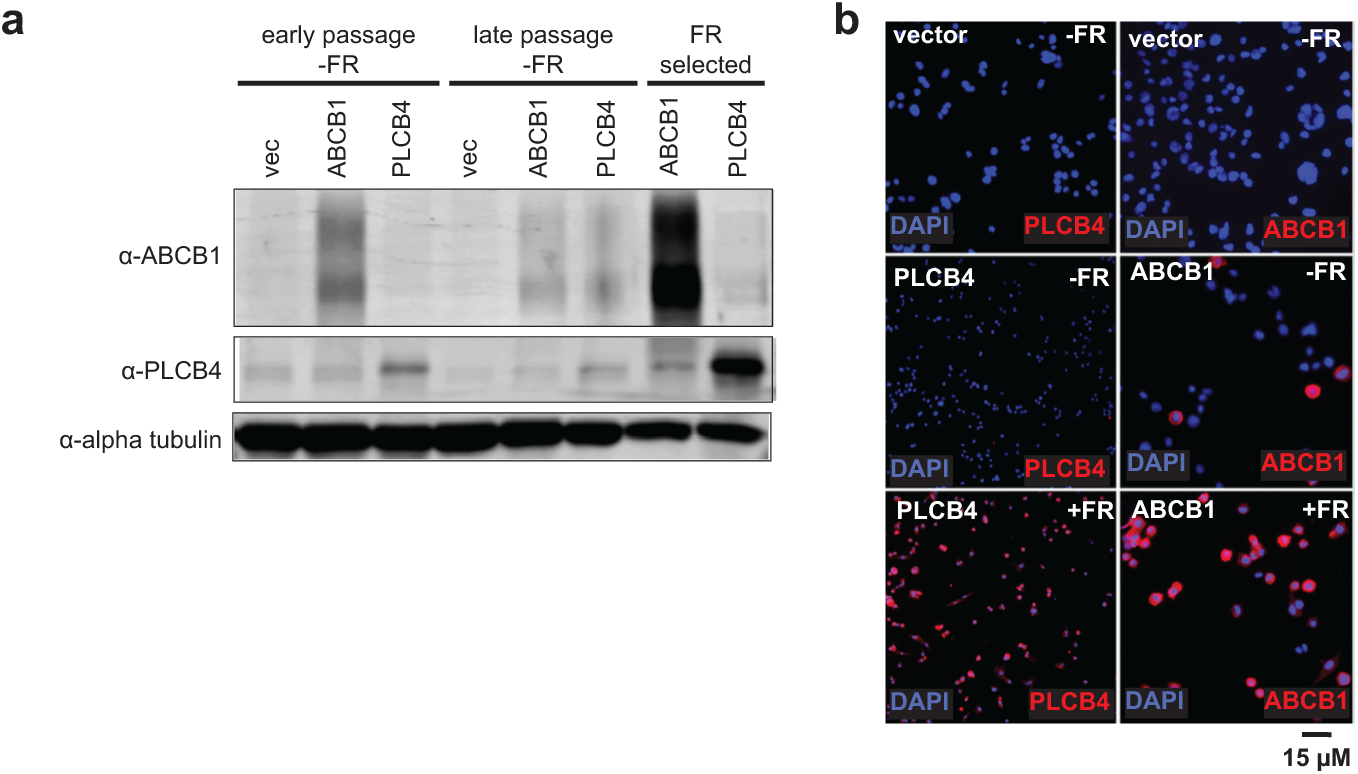
Transgenic expression of ABCB1 or PLCB4 is tolerated in a subpopulation of Mel202 cells. (**a**) Western blotting shows that both ABCB1 and PLCB4 are modestly expressed in cells prior to FR treatment. However, both proteins exhibit significant over expression in cells that have been selected on FR. (**b**) Immunofluorescence staining for either ABCB1 or PLCB4 shows that only rare cells over express the proteins prior to FR selection.

We performed immunofluorescence staining to evaluate protein expression in UM cells before and after FR selection (**Figure 4b**). Despite puromycin selection to ensure transgene expression, ABCB1 and PLCB4 protein was over-expressed in only rare UM cells. As expected, FR-resistant derivatives of these populations had more uniform over-expression (**Figure 4b**). We considered it unlikely that our transgene construct was undergoing frequent epigenetic silencing since we have routinely utilized the same vector in prior studies without observing a similar issue (Feddersen, Schillo, et al., 2019; Riordan et al., 2025; Schillo et al., 2024; Zhu et al., 2023). Instead, we suspected that over-expression of ABCB1 or PLCB4 was tolerated by rare UM cells in culture and that FR challenge selected for outgrowth of these cells.

We explored this possibility through additional experiments with ABCB1 given its relatively stronger performance as an FR-resistance driver (**Figure 2a,b**). A prior study expressed an ABCB1-GFP fusion protein and showed that the fusion protein maintains biological activity in cells (Wei et al., 2025). We generated a similar fusion protein by adding turboGFP to the C-terminus of ABCB1. This transgene was then cloned into a piggyBac transposon transgene along with a hygromycin resistance marker (EF1a:ABCB1-tGFP). A similar vector has also generated to express tGFP alone (EF1a:tGFP) or an empy vector (EF1a:empty). Each vector was then stably integrated into Mel202 cells. Following hygromycin selection, cells were analyzed by flow cytometry. As expected, EF1a:empty cells showed no evidence of tGFP expression (∼0.2%) while 98% of EF1a:tGFP cells showed strong tGFP expression (**Figure 5a**). Consistent with our immunofluorescence data, EF1a:ABCB1-tGFP cells showed significantly lower levels of tGFP (∼13%)(**Figure 5a,b**). The strong tGFP expression shows that the transgene construct is expressed well in Mel202 cells. Nevertheless, ABCB1-tGFP expression is not well-tolerated in most UM cells prior to sorting (**Figure 5b**). However, sorted Mel202 cells expressing high levels of ABCB1-tGFP maintain expression of the fusion protein (**Figure 5b**).

**Figure 5.**
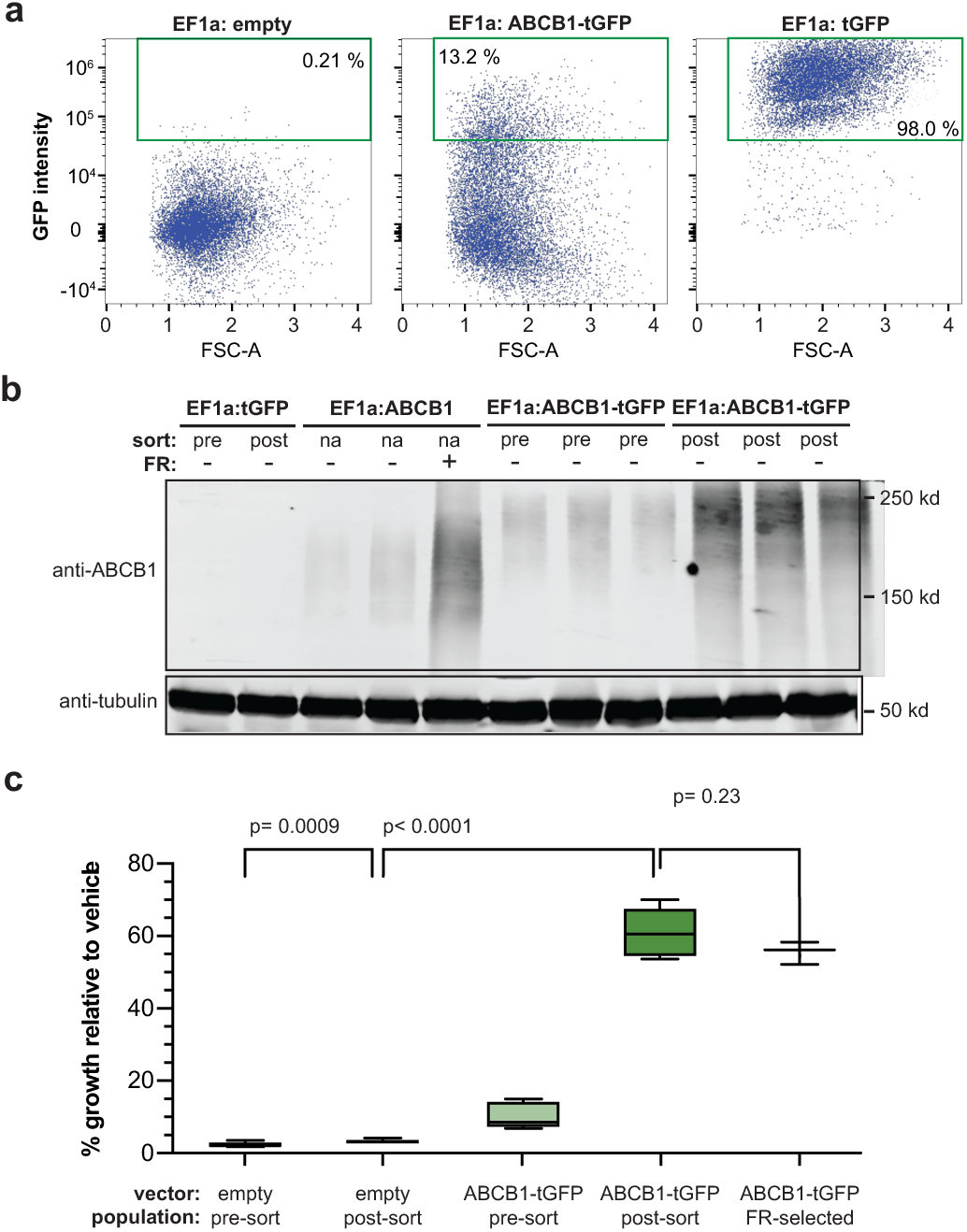
Expression of ABCB1-tGFP facilitates the isolation of FR-naïve Mel202 cells. **(a)** Following hygromycin selection, the majority of tGFP cells show strong fluorescence. By contrast, only 13% of ABCB1-tGFP cells have detectable fluorescence. **(b)** Western blotting confirms the expression of ABCB1-tGFP in sorted cells. **(c)** Isolated ABCB1-tGFP+ cells exhibit FR resistance that is indistinguishable from ABCB1-tGFP cells that were selected by chronic FR treatment.

We next utilized cell sorting to obtain a population of Mel202 cells that expressed high levels of ABCB1-tGFP. Importantly, these cells were FR naïve allowing us to then directly determine FR resistance using a cell population with more uniform ABCB1 expression. We measured growth using a resazurin assay at the end of seven days in media containing FR or vehicle alone to (**Figure 5c**). Consistent with our prior observation, ABCB1-tGFP expression in Mel202 cells prior to sorting weakly drove growth in FR relative to negative control cells expressing an empty vector (p=0.0009). However, sorted ABCB1-tGFP cells exhibit robust growth in FR that was statistically indistinguishable from FR-resistant ABCB1-tGFP cells that had been previously obtained by selection in constant FR (p=0.23)(**Figure 5c**). This result is consistent with the hypothesis that ABCB1 expression drives FR resistance in a subpopulation of Mel202 cells that can tolerate its expression.

#### ABCB1 ATPase activity is required for FR resistance in UM cells

While the efflux pump activity of ABCB1 has been shown to drive resistance to a variety of drugs (Skinner et al., 2023), there is evidence that ABCB1 can inhibit apoptosis in through a mechanism that is independent of its efflux function (*i.e.*, ATP independent)(Tainton et al., 2004). A prior study identified a missense mutation in ABCB1 (E556Q) that abolished its efflux activity (Zolnerciks, Wooding, & Linton, 2007). We generated a transgene to express the ABCB1^E556Q^ variant in UM cells. Expression of the variant in Mel202 and 92.1 cells was confirmed by western blotting (**Supp. Figure 4a**). Expression of the ABCB1^E556Q^ variant was higher in FR-naïve cells, particularly 92.1, when compared to wild type ABCB1 (**Supp. Figure 4a**). We then evaluated these cells for FR resistance a long-term assay in which cells were passaged under a constant FR dose (**Supp. Figure 4b**). This assay revealed that expression of the ABCB1^E556Q^ variant fails to provide resistance to FR.

#### ABCB1 expression is tolerated in UM cells with a lower rate of protein translation

Our findings support the hypothesis that ABCB1 promotes FR resistance in a subpopulation of UM cells that can tolerate its expression. However, the biological characteristics of ABCB1-tolerant UM cells are unknown. Therefore, we performed transcriptome profiling of three independent FR-naïve populations Mel202 cells that express ABCB1-tGFP or a tGFP control transgene (**Figure 5**). Differential gene expression analysis was performed to identify genes that showed altered expression in isolated ABCB1-tGFP cells. This analysis identified 355 genes – 48 with significantly higher expression (including ABCB1) and 307 with decreased expression (**Figure 6a**). Next, we performed gene ontology enrichment analysis to identify molecular functions, cellular components, or biological processes that were consistently altered in ABCB1-tGFP cells. This revealed that ∼15% of differentially expressed genes in ABCB1-tGFP cells are associated with translation and ribosome structure (**Figure 6b**).

**Figure 6.**
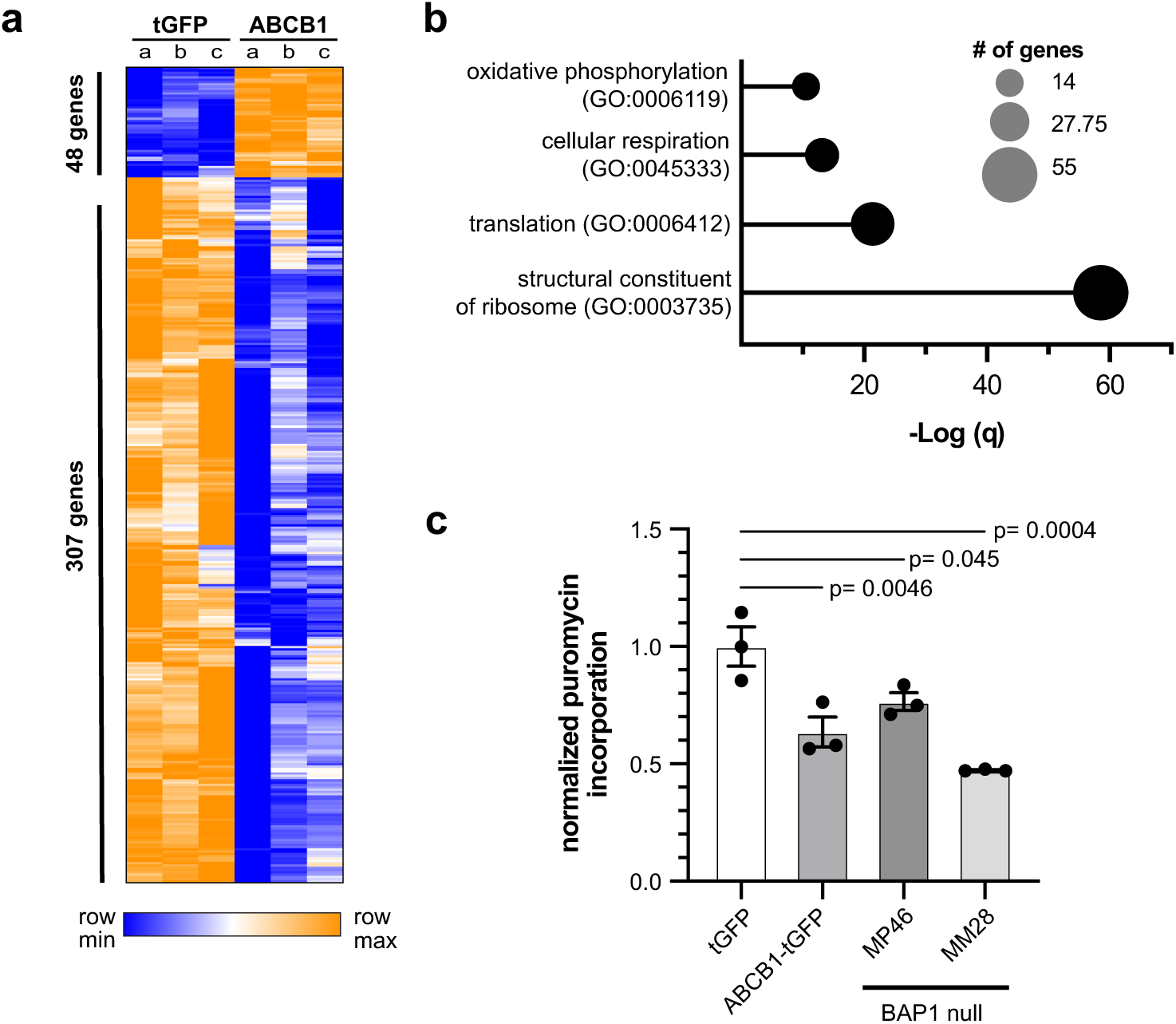
Mel202 cells that tolerate ABCB1 expression show decreased protein translation. **(a)** Transcriptome profiling of FR-naïve ABCB1-tGFP or control cells identified a set of 355 differentially expressed genes, the majority of which show decreased expression. **(b)** Gene Ontology enrichment analysis shows that ∼15% of differentially expressed genes are involved with translation or are structural components of the ribosome. **(c)** A puromycin incorporation assay determined the relative rates of translation in a panel of UM cells.

Several prior studies have implicated alterations in protein translation in various aspects of uveal melanoma biology (Chua et al., 2024; Dewaele et al., 2022; Johnson et al., 2017). Interestingly, Chua et al. showed that BAP1 loss in uveal melanoma cells is associated with reduced levels of pS6 leading to reduced proliferation and increased survival upon amino acid starvation (Chua et al., 2024). We directly quantified the rate of protein translation in UM cells using a puromycin incorporation assay (**Figure 6c**). This confirmed that ABCB1-tGFP cells do show a reduced rate of puromycin incorporation into proteins relative to control tGFP cells (p=0.0046). This suggests that the rate of protein translation in these cells is ∼60% of control cells and comparable to BAP1-deficient UM cells (MP46, MM28)(**Figure 6c**). However, unlike BAP1-deficient UM cells, Mel202 ABCB1 transgenic cells proliferate at a similar rate to control cells (**Figure 2a**).

## Discussion

We describe here a functional genomic screen using Sleeping Beauty transposon mutagenesis to identify drivers of resistance to FR900359 in uveal melanoma cell lines. Over expression of PLCB4 and ABCB1 was identified as the most frequent mechanisms of FR resistance to in our screens **(Figure 1).** However, predicted over expression of GRM1, LPAR3, and RGS4 were also identified as candidate resistance mechanisms. These genes provide an indication that alternative GPCR signaling may promote UM cell survival upon GNAQ/11 inhibition. GNAQ/11 signals to a multitude of downstream targets such as those within the MAPK pathway and we observed ERK, pS6, RasGRP3, and pP90 RSK S380 all had increased protein expression in UM cells overexpressed with ABCB1 or PLCB4 under FR treatment **(Figure 3**) (Chattopadhyay et al., 2016; Lapadula & Benovic, 2021; Park et al., 2018). The overexpression of ABCB1 or PLCB4 provides resistance to FR and maintains activation of the MAPK pathway oncogenic targets despite FR inhibition, compared to EV cells that had decreased protein expression levels of targets. These findings suggest that ABCB1 and PLCB4 contribute not only to FR resistance but allow continual activation of downstream MAPK targets associated with cell proliferation.

Our data suggest that the over expression of ABCB1 and PLCB4 drives FR resistance in a subpopulation of 92.1 and Mel202 cells **(Figure 2 & 4).** A proposed mechanism of acquired drug resistance is the induction of epigenetic modifications influencing expression of ABCB1, leading to a reduction in therapeutic drug response (Baker, Johnstone, Zalcberg, & El-Osta, 2005; Leschziner, Andrew, Pirmohamed, & Johnson, 2007). These observations implicate ABCB1 in cancer progression and drug resistance. ABCB1’s efflux pump has been shown to be the primary driver promoting drug resistance in ABCB1 cell population and altering the pump activity confirmed ABCB1’s efflux pump is necessary to promote resistance to GNAQ/11 inhibitor (**Supp. Figure 4**) (Hegedüs, Telbisz, Hegedus, Sarkadi, & Özvegy-Laczka, 2015; Schinkel & Jonker, 2003; Seelig, 1998; Szakacs, Paterson, Ludwig, Booth-Genthe, & Gottesman, 2006; Ueda, Cardarelli, Gottesman, & Pastan, 1987).

To define the cells that, creating a fluorescently tagged protein and generating a stable UM population that expresses ABCB1-tGFP and EF1α-tGFP, we observed by flow cytometry UM cells that express ABCB1-tGFP have a smaller proportion of cells that have increased tGFP fluorescence, indicative of ABCB1 expression compared to EF1α-tGFP UM cells with increased tGFP fluorescence (**Fig. 5A-C)**. This observation is consistent with our IF and immunoblot experiments in **Fig. 4C** where protein expression is low in overexpressed ABCB1 UM cells. This consistency provides evidence that UM cells do not tolerate ABCB1 expression well and the unique sub-population that could be possible markers for drug resistance and have high metastatic propensity (Landreville, Agapova, Kneass, Salesse, & Harbour, 2011).

ABCB1 overexpression has been shown to exhibit stem-like features (Balaji et al., 2020; Landreville et al., 2011). We hypothesize that there is a unique gene expression profile associated with UM cells that can tolerate ABCB1 over-expression. In order to identify the unique properties of this sub-set of cells that can tolerate ABCB1 expression without drug interference, we intend to determine gene expression profiles of ABCB1-tGFP sorted populations that are capable of sustaining ABCB1 high expression. Furthermore, we intend to evaluate ABCB1-tolerant UM tumor cell population in an in vivo model of disease progression as well as determine if increased ABCB1 expression predicts disease progression in UM patients.

Although FR has been used as a tool compound to inhibit GNAQ/11 due to its potent hypo-tensive effects, recently a GNAQ/11 inhibitor has been used in a first in human, phase I/II, multi-center clinical trial, NCT05415072, to treat metastatic UM patients. This trial uses DYP-688, a first-in-class PMEL17 melanoma-specific antibody-drug conjugate, as a single agent to deliver SDZ475 to inhibit GNAQ/11 oncogenic signaling resulting in dose-dependent apoptosis (Switzer, Piperno-Neumann, Lyon, Buchbinder, & Puzanov, 2023; van Dinten et al., 2005). Thus, using FR as a tool compound in our studies and understanding how UM tumor cells evolve resistance to GNAQ/11 inhibition is an important priority directly relevant to a clinical strategy targeting UM. We conclude that there are several genetic drivers that promote resistance to FR, but the two that have been validated here are PLCB4 and ABCB1. Both genes can be modestly overexpressed in UM cells but with FR treatment, the cells expressing these transgenes expand, creating a homogeneous population of PLCB4 or ABCB1 expression in UM cell populations. Future studies will be conducted to validate candidates from this SB screen such as performing single cell RNA-sequencing to determine distinct genetic profiles associated with FR treatment as well utilizing siRNA to knockdown candidates’ RNA expression to determine effects on FR treatment. Collectively, the data generated here may lead future discoveries to uncover tumor cell heterogeneity in UM and reveal a distinct subpopulation of UM tumor cells that is both capable of utilizing PLCB4 and ABCB1 to drive FR resistance.

## Acknowledgments

This work was supported in part by funding from the Department of Defense (W81XWH-22-1-0782) and the contributions of patients and families to the Iowa Melanoma Research Fund. Thank you to the generosity of primary investigators Kendall J. Blumer and Michael D. Onken from Washington University School of Medicine for gifting our UM cells, 92.1 and Mel202, as well as the therapeutic target, FR.

## Author contributions

S.D.M., J.D.R., C.S.S., and A.J.D. conceived the project. S.D.M. wrote the manuscript and performed most of the experiments. J.D.R. and A.J.D. performed the sequencing analysis. S.D.M and J.D.R. performed library preparation for UM SB screen. E.R.A. performed the western blots for GNAQ/11 downstream signaling and ABCB1-E556Q ABCB1 expression. J.D.R. performed growth assays for ABCB1-556Q. M.D.O. and K.J.B. provided the initial FR and UM cells. All authors contributed to manuscript revisions.

## Experimental model and study details

### Cell lines

All cells were grown in RPMI 1640 (Gibco) supplemented with 1% penicillin/streptomycin (Gibco 15140122) and 10% FBS (Bio-Techne # S11150). 92.1 and Mel202 were gifts from Drs. Blumer and Onken at Washington University School of Medicine. 92.1 has a mutation in EIF1AX, Mel202 harbors a mutation in SF3B1, and both 92.1 and Mel202 harbor the GNAQ mutation (Amirouchene-Angelozzi et al., 2014; Dahl et al., 2013; Dutton-Regester et al., 2012).

### Inhibitors

FR900359 (FR) was obtained as gift from Drs. Blummer and Onken at Washington University School of Medicine and are described in our experiments as FR-W. FR-W was used at a working stock of 50nM based on IC50 data **(suppl. Fig 2A).** Additional FR was purchased from Cayman Chemical (#3366) and is described in our experiments as FR-C. FR-C was used at a working stock of 5nM based on IC50 data **(suppl. Fig 2B).**

### *Sleeping Beauty* genetic screen

As previously described in other cell lines, 92.1 and Mel202 were stably transfected with SB100x using *piggyBac* (PB) transposon/transposase integration system with either Effectene (Qiagen) or jetOPTIMUS (Polyplus) (Feddersen, Wadsworth, et al., 2019; Wilson, Coates, & George, 2007; Yusa, Zhou, Li, Bradley, & Craig, 2011; Zhu et al., 2023). This system contains a PB transposase-expressing plasmid and an Ef1α-SB100X transgene embedded within a PB transposon (Feddersen, Wadsworth, et al., 2019). Following hygromycin selection, PB-Ef1α-SB100X positive cells were transfected with the pT2/Onc3 transposon plasmid, pooled together after cell confluency, and re-plated into ∼30 10 cm plates (Dupuy et al., 2009). Mutagenized (PB-Ef1α-SB100X-pT2/ONC3) and non-mutagenized (PB-SB100x-Ef1α) cells were treated with FR-W or vehicle (DMSO) 24 hours after seeding. FR was renewed every 7 days and vehicle treated cells were collected once confluent (∼3-4 days after initial drug dose) via trypsinization. Mutagenized cells treated with FR were collected upon colony formation, ∼21 or 28 days after initial treatment. DMSO and FR treated populations were harvested for DNA extraction. Sample and library preparations were developed with a stringent approach to determine transposon integration.

### Library preparation and sequencing

The identification of common transposon insertion sites across mutagenized resistant cells was ascertained via collection of genomic DNA from each plate using GenElute™ Mammalian Genome DNA miniprep Kit (Sigma). DNA fragments containing transposon/genome junctions were amplified via ligation-mediated PCR and sequenced using Illumina Hi-Seq 4000 platform (Feddersen, Schillo, et al., 2019; Riordan et al., 2014).

### Enforced expression of candidates

Cloned cDNAs of candidate genes were inserted into a piggyBac transposon expression vector and co-transfected with the piggyBac transposase in 10cm format (Zhu et al., 2023). The transfection workflow consisted of Qiagen Effectene (3ug DNA, 1:8 enhancer and 1:10 effectene ratios) for Mel202 and Jet OPTIMUS (5ug DNA and 5 μL reagent) for 92.1 (Zhu et al., 2023). Media was changed 24 hours after transfection for Mel202 and after 6 hours for 92.1. 48 hours post-transfection, cells were treated with hygromycin for ∼5 days at concentrations of 0.4 mg/mL for 92.1 and 0.2 mg/mL for Mel202.

### Viability Assays

Drug dose response curves using the resazurin assay were performed as previously described (Zhu et al., 2023). ∼5×103 cells were seeded in a 48-well plate, cells were treated with drug 24 hours after plating, and viability was measured in triplicate at each dose (Feddersen, Schillo, et al., 2019). IC50 data shown represents fluorescent signal detected at day 7 normalized to the day 0 reading acquired using the Biotek Synergy HT microplate reader and Gen5 software (Feddersen, Schillo, et al., 2019). Drug challenge data shown represents fluorescent signal detected at day 7 normalized to DMSO cell growth at day 7 using the Biotek Synergy HT microplate reader and Gen5 software (Feddersen, Schillo, et al., 2019). To generate a population doubling versus time (PDvT) long term growth assays, UM cells expressing PLCB4, ABCB1-wildtype, and ABCB1-E556Q were maintained in FR-W (50 nM), FR-C (5nM), or DMSO for ∼20-40 days. Every 5-7 days, UM cells were passaged, counted, and the population doubling level was calculated using the formula: PDLn = 3.32 (log Xt–log X0) + PDLn-1 (with Xt = cell number at that point, X0 = cell number used as inoculum and PDLn-1 = population doubling level at the previous passage) (Vanneste et al., 2020). Cells were seeded at 1×104 after cell counting. FR and DMSO treatment were renewed at each passage (Vanneste et al., 2020).

### Immunoblotting

15-20 mg of whole-lysate protein from UM cells and Biorad Precision Plus Protein Standards were loaded into 7-11% acrylamide gels, running at 140 volts. Lysates were transferred overnight at 4°C at 18 volts onto a PVDF membrane (Millipore IPVH00010). Western blot analysis was performed on an Odyssey infrared imaging system (LI-COR Biosciences) using Invitrogen goat anti-rabbit 790nM (A11369) and goat-anti mouse 680nM (A21058) fluorophores conjugated to: ABCB1 (C.S. E1Y7B), PLCB4 (sc-166131), pERK T202/Y204, (C.S 4370S), ERK (C.S 4696S), P90RSK S380 (C.S 11989S), phospho-P90RSK (Ser380) (D3H11) (C.S 119895S), phospho-S6 S240/244 (C.S 5364S), S6 (C.S 2317S), RasGRP3 (C33A3) (3334S), phospho-RasGRP3 (T133) (ab124823), and alpha tubulin (DSHB 12G10) primary antibodies. Technical duplicates were performed to confirm expression levels.

### Immunofluorescence Staining

Immunofluorescence staining (IF) was performed by carefully placing glass coverslips inside a 24-well plate, plating ∼1×105 cells/well in 1.5ml RPMI media, and allowing 24 hours for cells to adhere to the coverslip. Next, media was removed, and cells were fixed with 500ml/well of 10% formalin and incubated at room temperature for 15 minutes. Fixation was removed and cells were washed with 1XTBS 3x times. A blocking agent containing 10% goat serum plus NP40 was added to each coverslip and incubated at room temperature for 30 minutes. The ABCB1(C.S. E1Y7B) and PLCB4 (sc-166131) primary antibodies were added at 1:400 in 10% goat serum and incubated overnight at 4oC. Coverslips were washed with 1XTBS 3x times to remove excess primary antibody, the secondary antibody (Alexa Fluor 488-594, thermos fisher, A-11012) was added 1:400 in 10% goat serum, and incubated for 1 hour in darkness. Lastly, coverslips were washed with 1XTBS 3x times, washed with ddH2O, lightly dried with a chem wipe to remove excess liquid, and mounted cell side down to a glass slide using ProLong Diamond Antifade Mountant with DAPI (T.F. P36966). 20x images were taken using Lecia DMIRE2 Inverted Microscope.

### Flow Cytometry

UM cell populations expressing PB-ABCB1-turboGFP-PGKhygro (ABCB1-tGFP) and PB-EF1⍺-turboGFP-PGKhygro (EF1⍺-tGFP) were expanded until confluency in 10cm tissue culture plate. Upon day of analysis cells were trypsinized and cell number were counted using Thermo Scientific Invitrogen Countess 2 Automated Cell Counter. Pellets were resuspended in 1ml of ice cold FBS diluted in PBS, filtered in a 70μm cell strainer, and kept on ice until processing. Cells were sterile sorted using the Cytek Aurora Cell Sorter with 488 nm laser. Following sorting, cells seeded for expansion in RPMI media at 37°C.

---

**Supplementary Figure 1.**
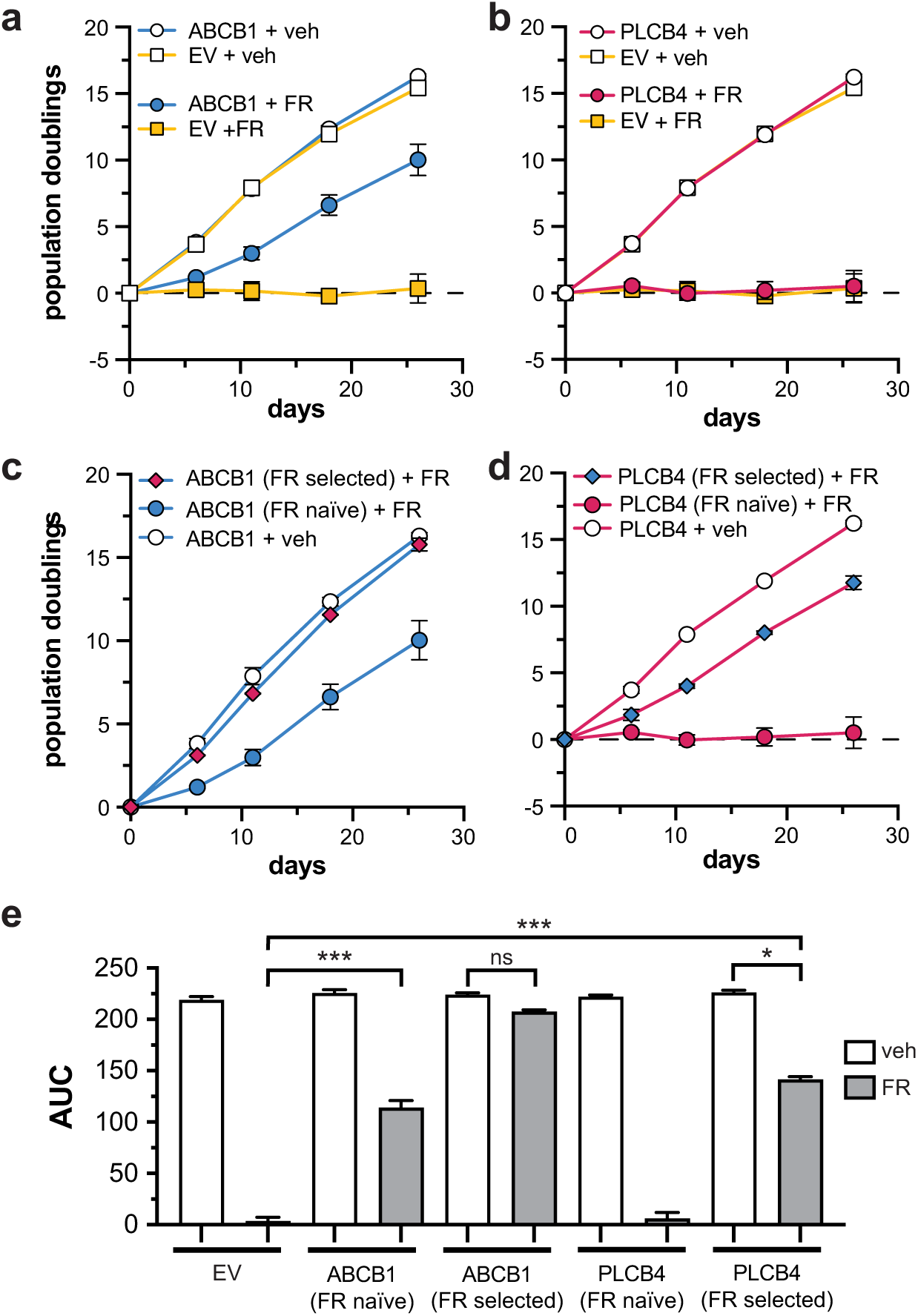
Validation of FR resistance mediated by over expression of ABCB1 and PLCB4 in Mel202 cells. Cell population doubling was monitored over several weeks in the presence of FR or vehicle for cells over expression ABCB1 (**a**) or PLCB4 (**b**). Over expression of either ABCB1 (**c**) or PLCB4 (**d**) led to the emergence of a cell populations that are indifferent to the presence of FR. (**e**) Quantification of the area under the curve (AUC) of the data shown in panels a-d.

**Supplementary Figure 2.**
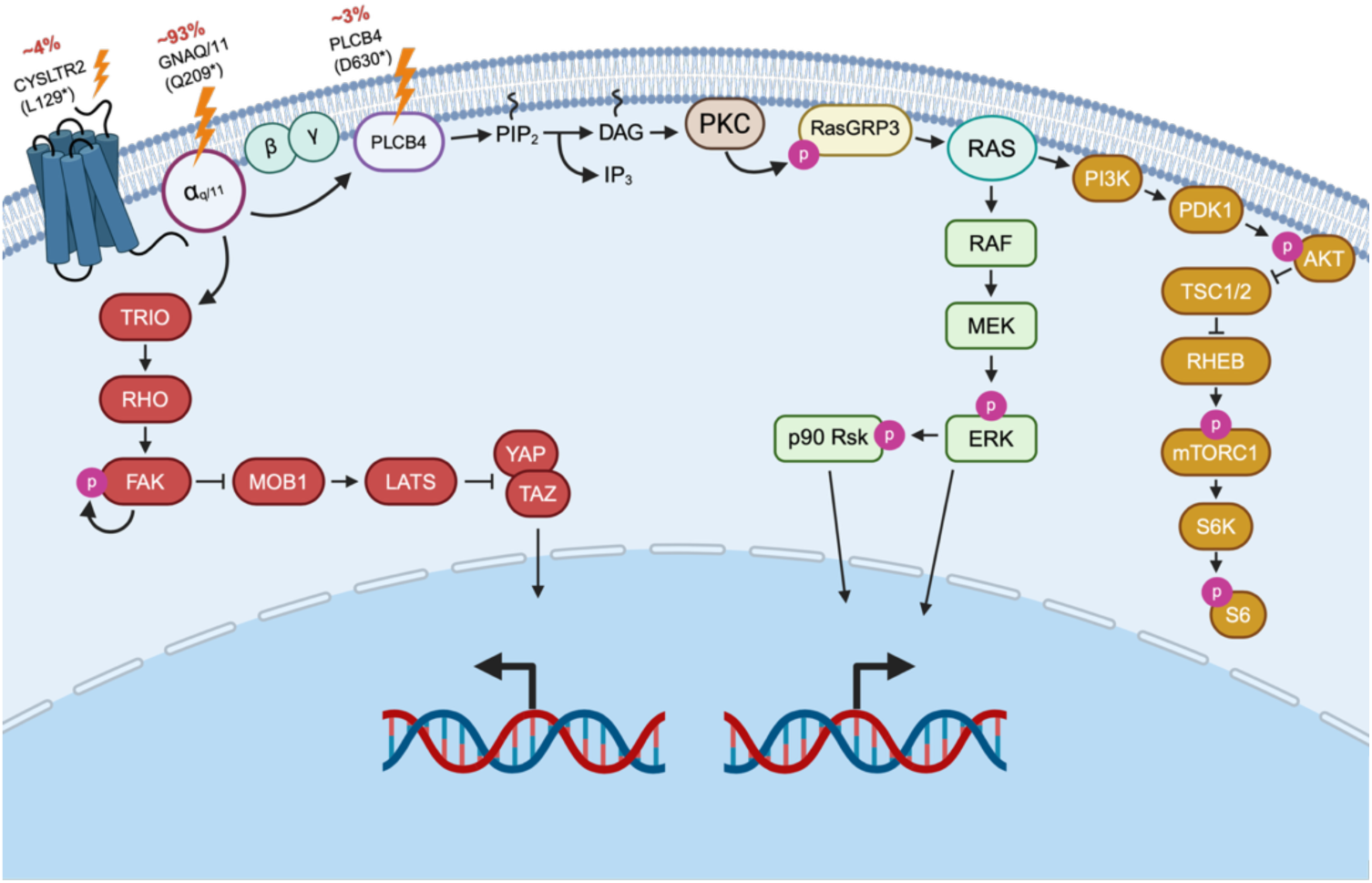
Overview of oncogenic signaling in uveal melanoma. Frequent somatic mutations in GNAQ, GNA11, PLCB4, and CYSLTR2 are mutually exclusive in most UM patients. These mutations are thought to mediate a complex signaling network involving the MAPK (green symbols), PI3K-AKT (orange symbols), and YAP/TAZ (red symbols) pathways.

**Supplementary Figure 3.**
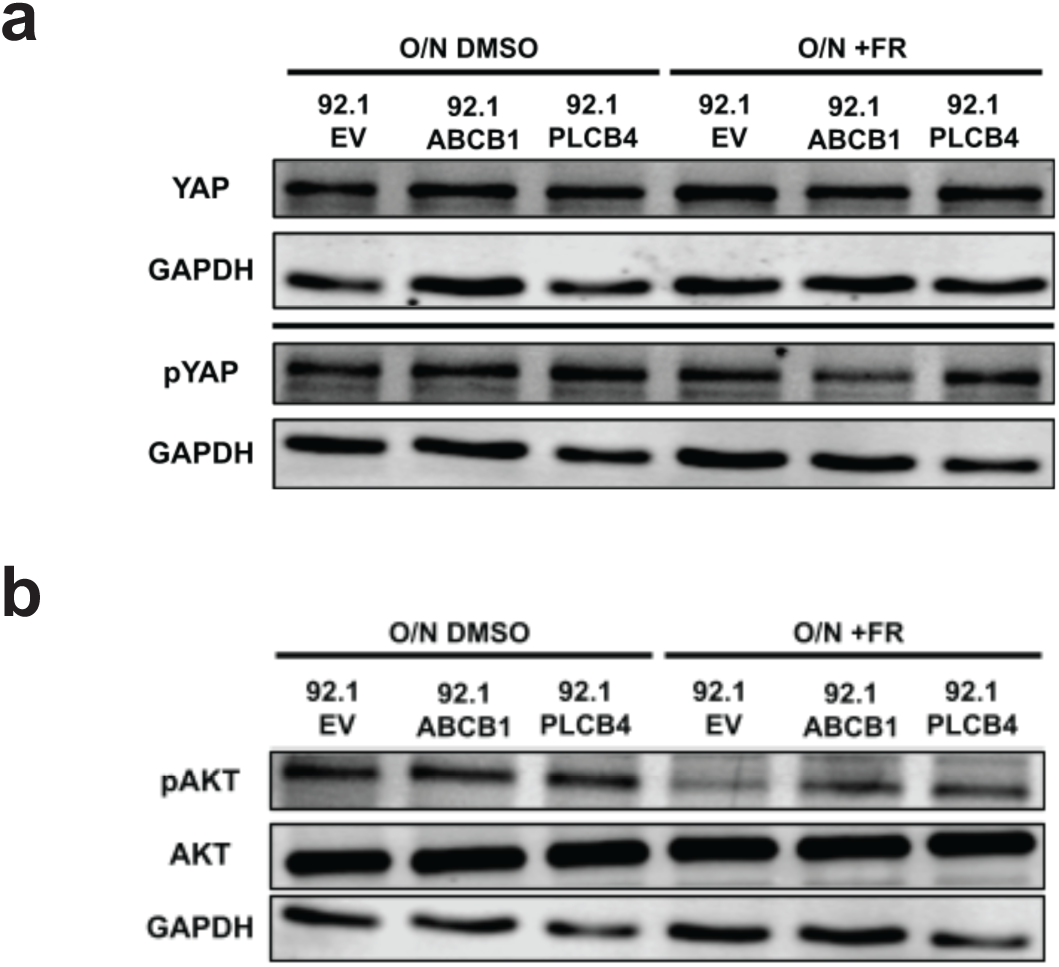
Effects of over expression of ABCB1 or PLCB4 on PI3K-AKT and YAP activity in 92.1 cells. **(a)** Levels of pYAP are unaltered by the over expression of either FR resistance driver. **(b)** The levels of pAKT are not as impacted by FR treatment in cells over expression ABCB1 or PLCB4.

**Supplementary Figure 4.**
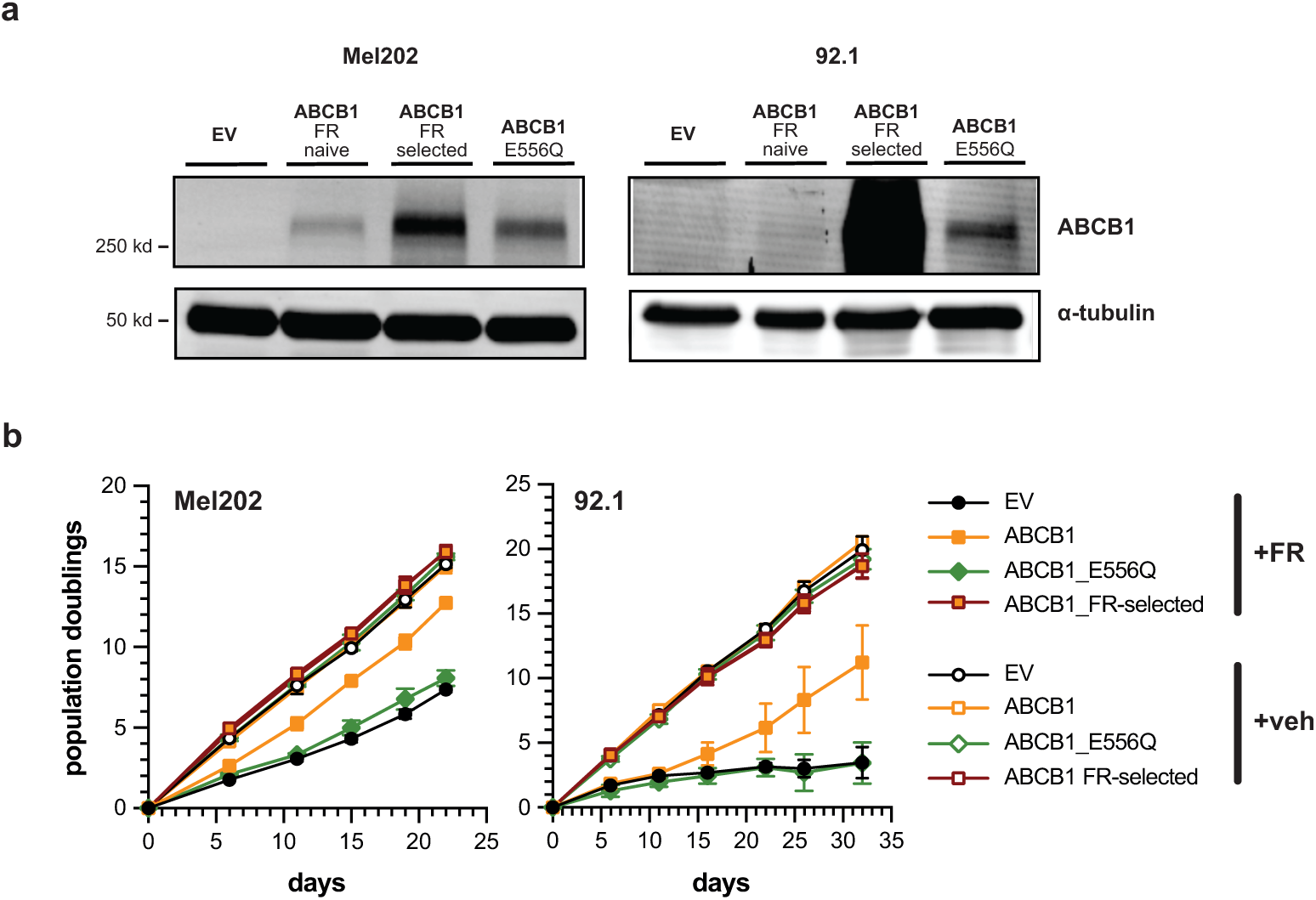
The ATPase activity of ABCB1 is required to drive FR resistance. **(a)** Expression of an ATPase deficient form of ABCB1 (E556Q) is better tolerated in both Mel202 and 92.1 cells compared to wild type ABCB1. **(b)** The inactive ABCB1 variant fails to drive FR resistance in culture.

